# Transcriptomic Insights into Early Diagnosis of Doxorubicin-Induced Cardiotoxicity in a Rat Model

**DOI:** 10.1101/2025.04.01.646673

**Authors:** Emilia Anca, Ilie-Ovidiu Pavel, Emilia Licărete, Corina Rosioru, Camelia Dobre, Manuela Banciu

## Abstract

Doxorubicin is a member of the anthracycline class of chemotherapeutic agents and is among the most effective treatments available up to date. However, its clinical use is significantly limited by severe cardiotoxic effects.

The purpose of this study is to investigate the transcriptomic alterations that occur in a rat model of doxorubicin-induced cardiotoxicity. Our results reveal significant dysregulations of cardiac metabolism and provide insights into the molecular mechanisms underlying cardiac damage produced by this treatment.

Our analysis revealed that heart tissue recovery following doxorubicin treatment is hindered by hypercholesterolemia exacerbated by transcriptomic-level alteration of the circadian rhythm. This result could help facilitate the discovery of circulating biomarker-based early diagnostic methods for this disease, along with cardioprotective treatment implementation.

## Introduction

Cancer continues to be a major burden globally as a significant obstacle to life expectancy, quality of life and health, in the swift transition to a rapidly aging population [1]. In 2022, it was estimated that the number of cancer cases was about 20 million worldwide, and by 2050, a 77% increase of these numbers is expected, bringing the anticipated cases of cancer to be over 35 million [2].

Despite the constant discovery and development of targeted molecular therapies for cancer treatment, which aim at reducing disease burden and long-term side effects, classic chemotherapeutic regimens continue to be a foundation in the treatment of most cancers due to their proven efficacy.

Anthracycline antibiotics are a class of chemotherapy drugs used in the treatment of numerous malignancies such as breast, ovarian, and colorectal cancers, along with Hodgkin and non-Hodgkin lymphomas, sarcomas, and leukemias [3]. The most known and used anthracyclines are daunorubicin, doxorubicin (DOX), epirubicin and idarubicin, which are included in approximately 60% of chemotherapy regiments administered to pediatric cancer patients [4]. The extensive use of anthracyclines is due to their great effectiveness [5]. Even though these are among the most used chemotherapeutic agents, their usage comes with a high price in the form of adverse effects that pose significant threats to the health and long-term well-being of cancer survivors [4].

The most concerning side effect of anthracycline use is pharmacological cardiotoxicity or cardiotoxicity (CTOX), which is characterized by the development of several heart-related conditions including arrhythmias, reduction in left ventricular ejection fraction, myocardial infarction and dilated cardiomyopathy. In the absence of early diagnosis and therapeutic intervention, these often progress to irreversible heart failure [6]. The precise pathogenesis of CTOX is not yet fully understood, however research shows that the increased oxidative stress produced by anthracyclines at heart level is most likely to be the cause [4,7].

Out of all anthracyclines, DOX has the most widespread use in clinical settings [8]. The present study, conducted in a rat model of DOX-induced CTOX, aims at consolidating current knowledge by confirming previously reported transcriptomic alterations identified in similar disease models.

The most known mechanisms by which DOX exhibits cytotoxic effects are related to direct DNA damage via topoisomerase 2 (TOP2) poisoning, overproduction of reactive oxygen species (ROS), and disrupting iron and calcium metabolism [5]. Despite the efficient antineoplastic effect, these mechanisms are also responsible for CTOX. A novel hypothesis, supported by recent research, DOX treatment accelerates the aging process in heart tissue. This accelerated aging, combined with the physiological changes associated to the normal aging – such as increased pro-inflammatory processes – contributes to heightened damage to the heart at multiple levels. These combined factors may worsen the long-term effects of DOX treatment, highlighting the need for further investigation into its impact on cardiovascular health [9].

Due to the complexity of this disease, finding early and non-invasive diagnostic techniques is the best way to combat pharmacological CTOX so that cardioprotective strategies can be implemented at the earliest sign of disease. In this study, we aimed to investigate the transcriptomic landscape of heart tissue in a rat model of DOX-induced CTOX and better understand which signaling and metabolic pathways are affected by the anthracycline treatment, with the final aim to associate these alterations with circulating biomarkers that could facilitate early diagnosis of CTOX.

## Materials and Methods

The present study adhered to the requirements of the European Directive 2010/63/EU, and national legislation 43/2014. The study protocol was approved by the Committee on the Ethics of Animal Experiments of Babe□ - Bolyai University, with the registration number 14.172/02.11.2021.

In our experiment, 6 adult male Wistar rats were divided into two groups. The DOX group (n=3) was treated with 3.75 mg/kg doxorubicin hydrochloride (European Pharmacopeia Reference Standard, D2975000, Merck administered weekly via tail vein injection for 4 weeks. The dosage was chosen to reach a cumulative dose of 15 mg/kg at the end of treatment, which is known to induce CTOX [10–12]. The Control group (n=3) received vehicle only (0.9% sodium chloride, B.Braun) in the same manner. The *i*.*v*. delivery of the therapeutic agent in the tail vein was chosen to minimize pain and inflammation which are known to affect animal welfare.

Heart tissue was collected and immediately placed in RNAlater Solution (Invitrogen), stored at 4°C overnight and then at −80°C until RNA was isolated according to instructions provided by the manufacturer. Total RNA was isolated using TRIzol Reagent (Invitrogen) and the RNeasy Mini Kit (QIAGEN) using manufacturer’s instructions. RNA was quantified using NanoDrop (Thermo Fisher Scientific).

RNA-Seq was performed by Macrogen (Macrogen Amsterdam, The Netherlands). Libraries were prepared according to the standard protocol for the Watchmaker mRNA library (Watchmaker Genomics, Inc.) and sequenced on the Illumina NovaSeq6000 platform (Illumina, Inc.).

The raw Fastq files were initially subjected to quality assessment via FastQC [13], followed by trimming and filtering using Trimmomatic [14] to eliminate low-quality bases and adapter sequences. The trimming parameters were as follows: ILLUMINACLIP:2:30:10, LEADING:10, TRAILING:10, SLIDINGWINDOW:4:15, HEADCROP:12, CROP:85 and MINLEN:36. The processed reads were then aligned using the STAR aligner [15]. To generate the genome index, the reference genome Rattus norvegicus mRatBN7.2 (https://www.ncbi.nlm.nih.gov/datasets/genome/GCF_015227675.2) FASTA file and the corresponding RefGene GTF annotation from UCSC were utilized [16]. Gene-level quantification was subsequently performed with HTSeq [17], counting reads that mapped to exons (-t exon), associating counts with gene identifiers (-i gene_id), and resolving read overlaps using the union model (-m union).

Differential gene expression analysis was performed using DESeq2 [18]. Enrichment analyses were conducted using Gene Ontology [19,20], Gene Set Enrichment Analysis [21] and the Kyoto Encyclopedia of Genes and Genomes [22], and data visualization was executed using the Pathview R package [23]. Statistical significance was considered at p-values <0.05.

The data discussed in this publication have been deposited in NCBI’s Gene Expression Omnibus (GEO) and are compliant with the MINSEQUE (Minimal Information About a Next-generation Sequencing Experiment) and are available under the GEO Series accession number GSE288128 (https://www.ncbi.nlm.nih.gov/geo/query/acc.cgi?acc=GSE288168).

## Results

### Overall doxorubicin effects on the gene expression in the cardiac tissue

The principal component analysis (PCA) plot of counts, after DESeq2 and rlog normalization, showed clustering between the samples collected from the Control and DOX groups (Figure 1A). This result shows that the transcriptional landscape of heart tissue is altered following DOX treatment.

**Figure 1.**
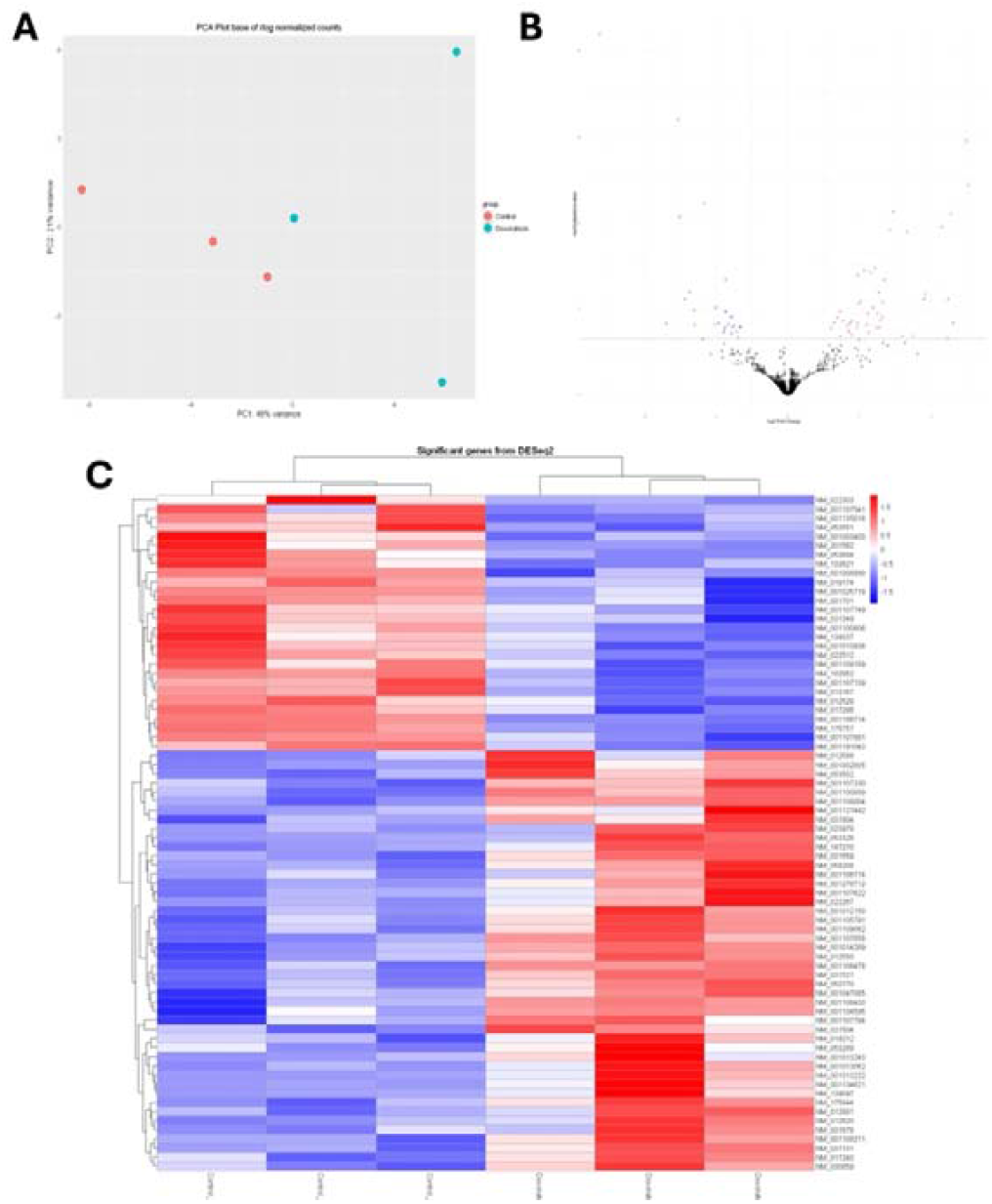
Transcriptomic analysis of heart tissue from DOX-treated (n=3) and control (n=3) rats. A. PCA plot showing clustering of the samples; B. volcano plot of differential gene expression, with log2 fold-change (x-axis) versus log10 adjusted p-value (y-axis), and significant dysregulated genes highlighted in blue (downregulated) and red (upregulated); C. heatmap of top differentially expressed genes with hierarchical clustering applied to both genes and samples.

We then shrunk the results of DESeq2 to investigate the differentially expressed genes (DEGs) using ‘apeglm’ [24]. At an adjusted p-value of <0.05, we identified a total of 79 DEGs (Figure 1B) in our dataset, out of which 28 were downregulated and 49 were upregulated. Among upregulated genes, we identified actin-binding genes (Acta1 and Shroom3), Adamts19 involved in collagen fibril organization, the transmembrane transporter Slc26a1, Ccnd2 associated with cell survival pathways and the transcription factor Hltf. In the list of downregulated genes, we identified genes associated with stress responses such as Pitpnm2, Card9, Macrod1 and Tp53i11, and several others also involved in cell signaling: Aplnr, Cxcl11, Lingo4 and Pxdc1 (Figure 1C and Supplementary Table 1).

### Signaling pathway analysis in cardiac tissue after cumulative exposure to DOX

Next, we sought the pathways and processes associated with the discovered DEGs. We performed gene set enrichment analysis (GSEA) to discover the biological pathways altered by DOX treatment. Using the list of DEGs, we identified that pathways involved in transcription (GO:0006351, 9 genes - Tgfb2, Creb5, Spp1, Sfrp1, Ankrd23, Nr1d2, Mdk, Abra and Per2), and RNA metabolic processes (GO:0016070, 13 genes - Tgfb2, Creb5, Per3, Spp1, Sfrp1, Ankrd23, Nr1d2, Mdk, Abra, Bhlhe40, Irx4, Per2, and Lrrfip1) are significantly enriched following DOX treatment, while cell junction assembly (GO:0007043, 4 genes - Fscn1, Aplnr, Cldn5, Hopx) and Lingo4) and organelle membrane (GO:0031090, 7 genes – Rtn2, Tmem86a, Acads, Car4, Cyp11a1, Ryr2, and Ucp3) processes were suppressed (Figure 2A).

**Figure 2.**
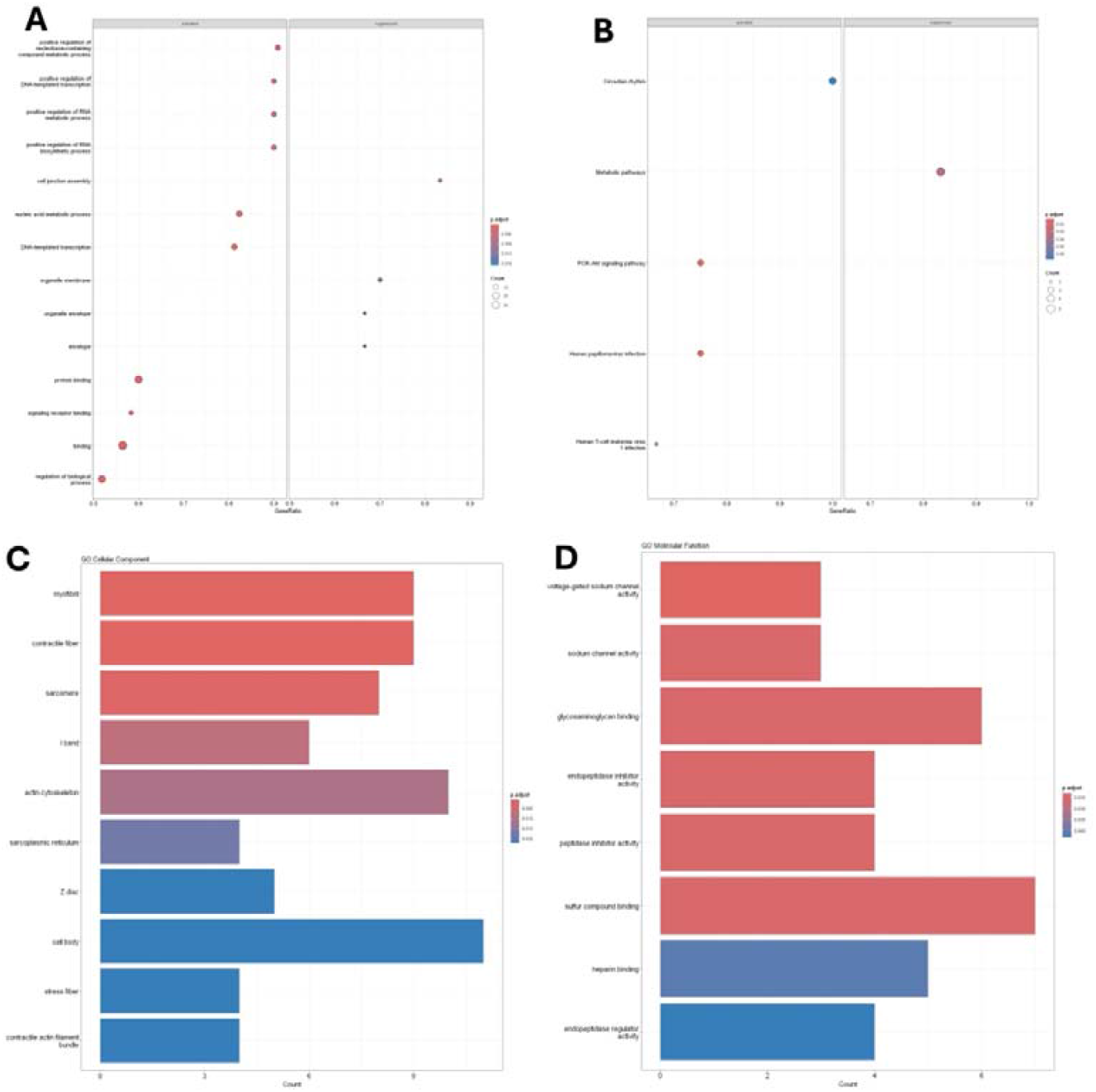
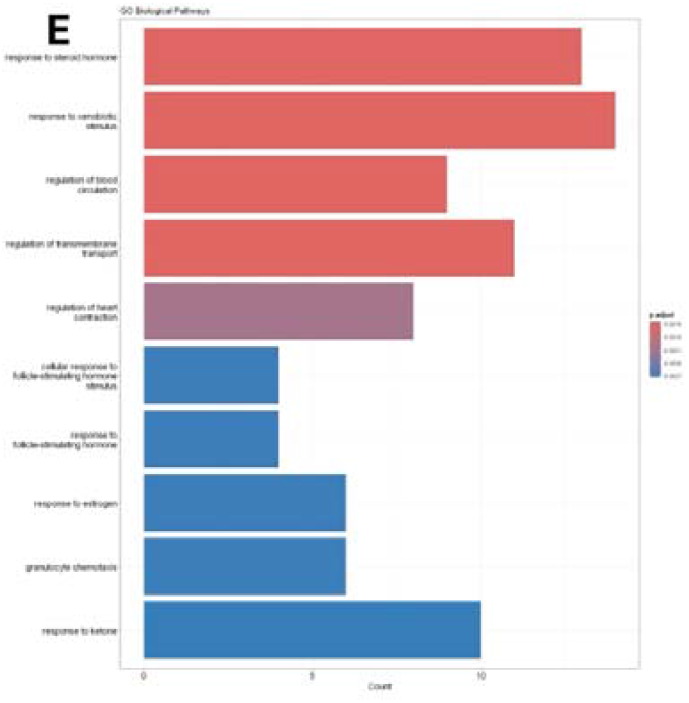
Pathway analysis of DEGs in heart tissue post-DOX treatment. A. GSEA of activated and suppressed pathways in the DOX-treated group (n=3) compared to Control (n=3); B. pathways identified by KEGG; C. analysis of GO biological functions, D. GO molecular functions, and E. GO cellular component, sorted by p-adjusted value.

Gene Ontology (GO) of cellular components showed that myofibril, contractile fiber, and the sarcomere are the most affected components following DOX treatment (Figure 2C). Between molecular functions, enriched terms in our comparison are related to sodium channel activity (GO:0005248, GO:0005272), glycosaminoglycan binding (GO:0005539) and pepdidase inhibitor activity (GO:0030414, GO:0004866) (Fig. 2D). For biological processes, regulation of blood circulation (GO:1903522, 9 genes - Tgfb2, Scn3b, Myh7, Per2/, Myh7b, Ednra, Hopx, Scn4b, Ryr2), cardiac muscle contraction (GO:0008016, 8 genes - Tgfb2, Scn3b, Myh7, Myh7b, Ednra, Hopx, Scn4b, Ryr2), and response to steroid hormone (GO:0048545, 13 genes - Tgfb2, Spp1, Sfrp1, Mdk, Ccnd2, Serpine1, Acta1, Hcn2, Acads, Card9, Car4, Cyp11a1, Ucp3) were significantly altered by DOX treatment (Fig. 2E).

The Kyoto Encyclopedia of Gene and Genomes (KEGG) enrichment analysis identified that DOX treatment led to the activation of circadian rhythm (Fig. 3A), phosphatidylinositol 3-kinase – protein kinase B (PI3K-Akt) signaling and adrenergic signaling (Fig. 2B, and Fig.3BC) were activated while suppressed genes were generally associated with the general ‘metabolic pathways’ term (Fig. 2B).

**Figure 3.**
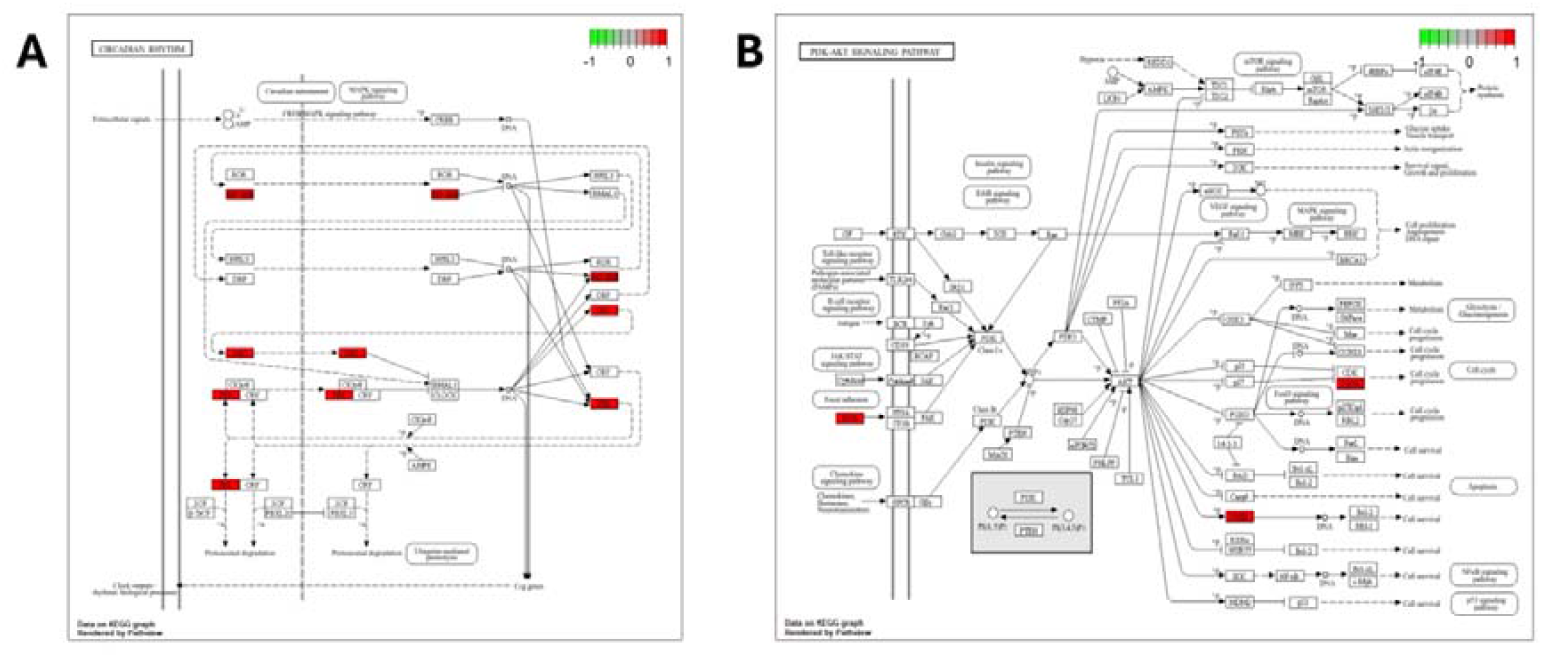
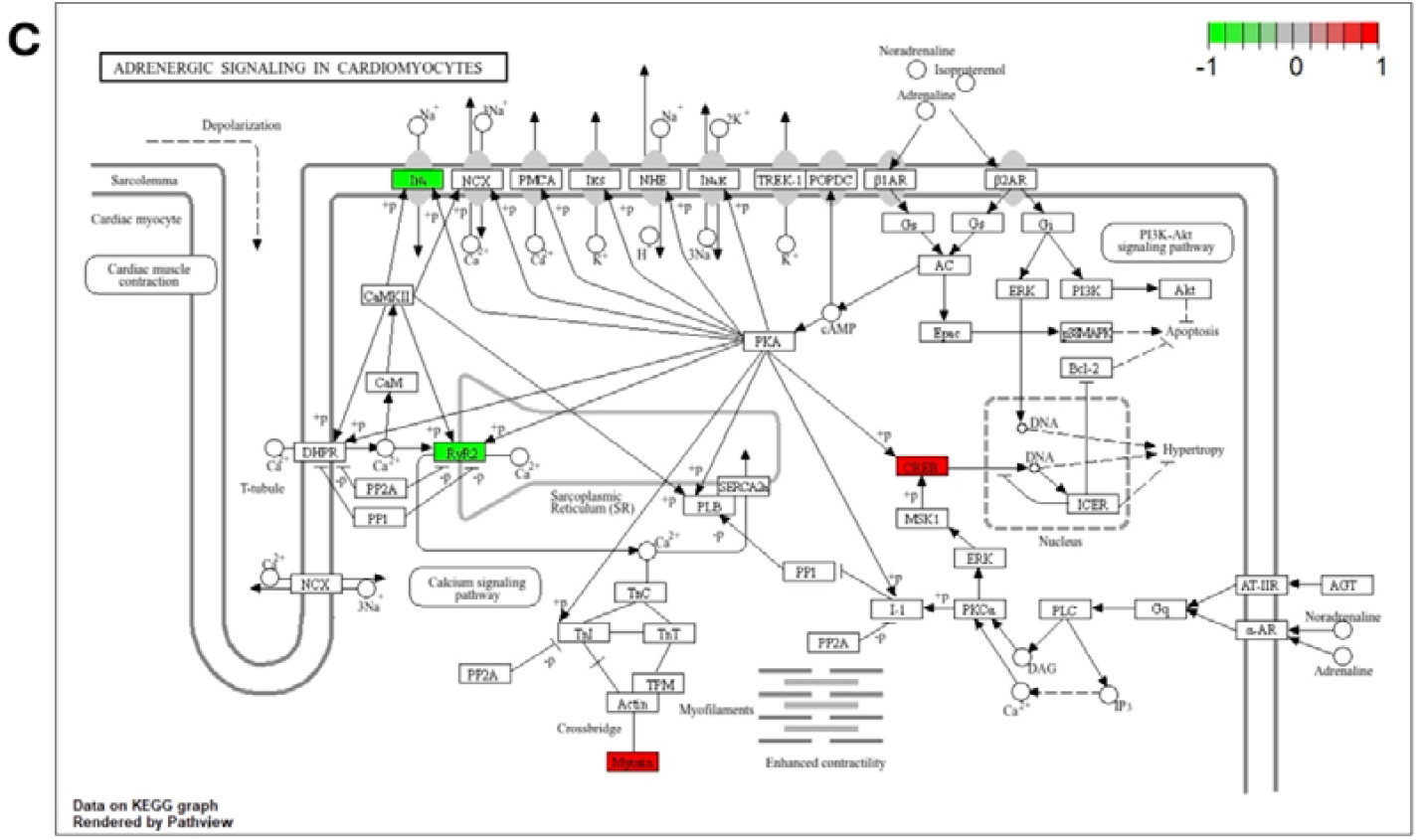
KEGG pathway analysis of rat tissue following DOX treatment. A. Circadian rhythm; B. The PI3K-AKT signaling pathway; C. Adrenergic signaling in cardiomyocites. ECM – extracellular matrix; CREB - cyclic adenosine monophosphatase response element binding protein; REV-ERB - nuclear receptor subfamily 1, group D, member 1; PER – period circadian clock; DEC - D-box binding protein

## Discussion

This is a follow-up study of our recent data which showed that CTOX effects induced by DOX administration could be identified shortly after the completion of treatment by evaluating a battery of circulating biomarkers [25]. Thus, we noted that fulminant CTOX-inducing effects of DOX tend to subside slowly in the post-treatment period (hypercalcemia and iron overload), while some are protracted and pose long-term risks for cardiovascular health (hyperlipidemia).

To gain insight the mechanisms that support our recent findings we performed transcriptome profiling of heart tissue from rats subjected to a DOX regimen known to induce CTOX, as demonstrated by Podyacheva et al., 2021 [12]. These data could ensure a step forward into the identification of biomarkers for cardiac damage associated with DOX administration in cancer patients for early non-invasive diagnosis.

To this aim, we investigated the transcriptomic landscape of heart tissue in a rat model of DOX-induced CTOX. We first evaluated DEGs between DOX- and vehicle-treated samples, and performed functional enrichment analysis through GO, GSEA and KEGG.

The PI3K-Akt pathway is a key intracellular signal transduction pathway, with critical role in maintaining heart tissue homeostasis by controlling many processes such as cardiomyocyte growth, survival and stress responses [26]. Moreover, it is a key player in the development of cardiac disease, by mediating cardiomyocyte injury following oxidative stress and is a discovered therapeutic target in DOX-induced CTOX [27,28].

In our experiment, using KEGG we identified upregulated members of the PI3K-Akt pathway - ECM and the cyclic adenosine monophosphatase (cAMP) response element binding protein (CREB), most likely as a result of fibrosis [29,30] induced by DOX administration. As cardiomyocytes are terminally differentiated shortly after birth [31], the upregulation of Cyclin could indicate fibroblast proliferation which are activated following tissue injury [32]. This result can be corelated with our previous finding regarding an increase in plasma levels of NT-proBNP levels following DOX administration [25], which is a known marker of cardiac tissue fibrosis [33].

We also identified the impairment of the adrenergic signaling pathway, which plays important roles in contractility regulation [34]. The major metabolite of DOX, doxorubicinol (DOXol) disrupts the normal Ca^2+^ handling directly by binding to the ryanodine receptor 2 (RYR2) or indirectly by reactive oxygen species (ROS) overproduction. This is a known contributing factor to CTOX [35]. RYR2 is the main channel responsible for calcium release from the sarcoplasmic reticulum, and a decrease in its activity leads to impaired contractility, which is one of the major causes for arrhythmias [36]. The decreased activity of RYR2, coupled with the upregulation of myosin, suggests an attempt to counteract the increased strain on the cardiomyocyte contractile system following DOX administration. This result is consistent with our previous work, where we observed a higher plasma calcium concentration after following the same DOX administration protocol [25].

Lastly, we identified an intense disturbance of circadian rhythm (CR) molecular components by the enrichment of nuclear receptor subfamily 1, group D, member 1 (REV-ERB), D-box binding protein (DEC), and period circadian clock (PER) genes, which all have other non-circadian functions. The normal function of the entire cardiovascular system heavily depends on the CR [37] and disturbances in this rhythm impairs heart function, thus contributing to oxidative stress inflammation and cardiovascular disease. REV-ERBs α and β contribute to the fine-tuning of glycerolipid metabolism in heart tissue [38]. Specifically, upregulation of these receptors leads to a decrease in cholesterol efflux and reduced lipogenesis, thus lowering lipid concentration in plasma [39]. In our previous study, we identified that DOX treatment is associated time-dependent increases in cholesterol and triglyceride plasma levels [25]. Taken together, these results suggest that DOX-induced dyslipidemia could be attributed to liver dysfunction and not on REV-ERB impairment.

PER genes control the DNA response as result of damage apart from regulating the CLOCK-BMAL1 transcription factors [40]. The DEC transcription factor activation leads to the reset of the clock, due to its role as a negative regulator of CLOCK-BMAL1 [41]. The disruption of the circadian rhythm was also seen in mice treated with DOX [42], and a direct link between DOX administration and impaired sleep was also found in rats [43]. Sleep disturbance is a known contributor to hypercholesterolemia and increased lipid levels in the liver - an organ that is already severely impacted by DOX pharmacokinetics [44,45].

A major limitation of the present study is the small sample size, which may have affected the interpretation of our results despite several methods of analysis. Additionally, we only focused on transcriptomic profiling, without validating the obtained data. Despite these constraints, our results are consistent with those observed in similar studies, thus strengthening the validity of our results. Future studies, integrating multiomic analyses with larger sample sizes and other tissues known to be affected by the toxicity of DOX are crucial for a comprehensive understanding of the underlying processes affected by this treatment.

## Conclusion

Our study aimed to characterize the transcriptomic alterations that occur in cardiac tissue in a rat model of DOX-induced CTOX. Although limited by the small number of samples, these findings offer a valuable overview of the molecular mechanisms disrupted by DOX. We observed significant dysregulation of pathways involved in inflammation, hormone signaling and circadian rhythm regulation, which are all known to contribute to the deleterious side effects of DOX on cardiac function. Moreover, we were able to link present transcriptomic findings with our previous assessment of circulating biomarkers in the same CTOX model.

In addition to the hyperlipidemic effect of DOX observed in our previous study, our transcriptomic analysis revealed significance alteration of genes associated with the regulation of circadian rhythm. As disruption of the circadian rhythm is a known exacerbator of hypercholesterolemia, which is linked to cardiovascular disease pathogenesis, our results may indicate that DOX-induced circadian rhythm disturbance may contribute to the worsening of lipid metabolism impairment and further sustain the CTOX-promoting environment in the heart, which could prevent or delay cardiac tissue recovery following DOX treatment.

These results highlight the importance of further investigations to develop a deeper understanding of DOX-induced CTOX, which is essential for identifying biomarkers that enhance early diagnosis capabilities, as well as developing better therapeutic strategies to prevent cardiac damage associated to this treatment, which is still widely used in treating numerous types of cancer.

